# Gestural Communication of Wild Bonnet Macaques in the Bandipur National Park, Southern India

**DOI:** 10.1101/739441

**Authors:** Shreejata Gupta, Anindya Sinha

## Abstract

Nonhuman primate gestures are believed to be crucial evolutionary precursors of human language. Comparative studies on primate gestures in an evolutionary framework have, however, remained largely restricted to the great apes and the potential flexibility and richness of gestural communication in monkeys, especially in the wild, continue to be virtually unknown. In this paper, we followed several criteria, adapted from ape gesture studies, to identify gestures and evaluate their contexts of usage in the repertoire of wild bonnet macaques Macaca radiata in the Bandipur National Park of southern India. This report is the first of its kind to systematically identify gestures in any wild, non-ape species, thus providing a platform for comparative studies across primate taxa, particularly in our efforts to trace out the phylogenetic origins of language-like markers in the primate lineage, earlier than in the great apes.

## Introduction

Investigations into nonhuman primate gestures initially employed traditional methods largely derived from studies on human gestures (Pika 2008; Scott and Pika 2012; Tomasello et al. 1994; Tomasello et al. 1997; Tomasello et al. 1985; Pika, Liebal, and Tomasello 2003, 2005; Liebal, Pika, and Tomasello 2006, 2004). Gestures in nonhuman primate communication, especially those of apes, were thereafter defined as “discrete, mechanically ineffective, physical movements of the body” (Hobaiter and Byrne 2011a; Genty et al. 2009; Cartmill and Byrne 2010) or “communicative signals produced by body postures or the movement of body parts, including the limbs, head and /or facial muscles to achieve an intended goal” (Tomasello and Call 2007). It has also been suggested that gestures should be signals always directed to a particular recipient and be mechanically ineffective in eliciting voluntary responses from the recipient (Arbib, Liebal, and Pika 2008; Pika 2008; Schel et al. 2013). Currently, summarising from the literature, a generally accepted operational definition of “gesture” (in the great ape literature) is: *a discrete, mechanically ineffective movement of body parts deployed intentionally (with the aim of eliciting a behavioural response from the recipient)*.

Like the study of tool use in nonhuman animals, research on great ape behaviour has frequently led to the discovery of similar behaviour in other primates. A recent review of ape gestures, compiling findings from the past six decades, supports the hypothesis that gestures constitute innate signals which can be used in intentional ways to convey particular meaning in communicative contexts (Byrne et al. 2017). As gesture forms appear phylogenetically continuous among apes, it is quite possible that similar gestural communication may be encountered in non-ape primate species as well. Unfortunately, earlier investigations of gestural communication in macaques did not use the same operational definitions of “gesture”, as stated above (Laidre 2012, 2008, 2011; Maestripieri 1996, 1997, 2005). Maestripieri (1996, 1997, 2005) defined gestures as a collective term for facial expressions, hand gestures as well as body postures. Laidre (2012, 2008, 2011) reported and examined novel gestures in captive mandrills, their developmental histories and social learning processes; Hesler and Fischer (2007), described gestural communication of barbary macaques in captivity, following operational definitions established in the primate gesture literature. In this paper, we revisit the natural behaviours documented from a population of wild bonnet macaques *Macaca radiata* in the Bandipur-Mudumalai forest ranges of southern India (Sinha et al. 2005; Sinha 2003, 2005, 2014; Dutta-Roy and Sinha 2001; Chatterjee 2013) and test and test whether macaque gestures follow the same operational definition described above.

First, to examine whether macaque gestural communication uses “discrete” signals, we considered *instantaneous* communicative behavioural events displayed by our study individuals as potential gestures and not those that prevailed over longer durations of time. Thus, we consider gestures to be behavioural events rather than behavioural states (Altmann 1974); this is in accordance with the ‘discrete’ nature of gestures suggested by previous studies (Genty et al. 2009; Cartmill and Byrne 2010; Hobaiter and Byrne 2011a). To distinguish the function of each potential gesture and to avoid the functional complexity of multiple gestures, we also considered only those instances where a single act was displayed by the signaller to initiate communication.

Second, the sent signals were “mechanically ineffective” (Arbib, Liebal, and Pika 2008; Pika 2008; Schel et al. 2013), meaning that they did not physically enforce the recipient to produce a response. The responses were thus necessarily voluntary. For example, a mother’s physically picking up of her eager infant would be considered a mechanically effective action but one that involves her extending her arms towards the child with upward-facing palms, resulting in the infant rushing to her, would be considered mechanically ineffective. The criterion of mechanical ineffectiveness does *not* exclude potential gestures that are tactile, i.e. gestures that involve the signaller coming into direct physical contact with the recipient. In such cases, a tactile gesture must have a response that is voluntarily executed by the recipient and not a mere physical reaction to the action. Thus, an accidental collision between two individuals resulting in one falling down is *not* a gesture-response pair; a ‘slap’ and an eventual movement of the receiver’s face is also *not* one; a ‘slap’ followed by a voluntary response that is not result of the physical impact of the signaller’s action, such as ‘fleeing’ or ‘lunging’, however, *is* a gesture-response pair.

Finally, we evaluated whether the potential gestures that we identified were used “intentionally” by bonnet macaques. A signal is defined to be intentional when an individual performs an act with the aim of changing the behaviour or mental state of another individual. Non-intentional communication, in contrast, occurs through reflexive signals produced in response to a stimulus, as, for instance, an immediate scream emitted on accidentally touching a hot surface. We thus consider communication that effectively changes the behaviour of another individual as a marker of intentional use of signals, irrespective of the underlying mental states of the signaller or the receiver, which is otherwise difficult to ascertain (see (Townsend et al. 2017)). Great ape gestures, in fact, have been considered to be intentionally produced whenever they are goal-directed (Plooij 1977; Piaget and Cook 1952; Bard 1992; Plooij 1979).

In order to objectively define the markers of intentional use of primate gestures, several criteria have been adapted from those laid out for human language studies (Piaget 1981; Bruner 1981; Bates, Camaioni, and Volterra 1975; Townsend et al. 2017; Pika, Liebal, and Tomasello 2003; Tomasello et al. 1994; Tomasello et al. 1985; Leavens and Hopkins 1998). According to these established criteria, the signals should be (i) socially used, targetted to a recipient; (ii) adjusted to the recipient’s body orientation; (iii) followed by successive visual orientation or gaze alternation between an object of interest or event; (iv) accompanied by attention-seeking tactics of the signaller; (v) followed by a persistent display of the same or other signals, in the absence of an initial response; and (vi) elaborated further, in the absence of the initial response. The first four criteria are crucial to establish the premise for intentional signaling while the last two are testimony to goal-directedness, thus signifying the purpose of such intentional behaviour.

In this analysis, we considered only those interactions in which the signaller was directly looking at a target recipient—a necessity in order to produce intentional behaviour—and a subsequent response was elicited (Tomasello et al. 1994). There were, however, occasions when there was visual contact established between the signaller and the receiver but the latter moved away from the signaller. We have not considered these interactions here, as the motivations behind such responses were difficult to interpret and did not reflect the communication context in which the signaller initiated the interaction. In certain other circumstances, there was an initiation of a potential communication event, involving at least two individuals, but the receiver did not visually attend to the act; such instances have also not been considered in this analysis.

This paper, therefore, documents our search for gestures in the behavioural repertoire of wild bonnet macaques. We explore the outcomes of interactions in which the signaller directed a signal to another individual, who was visually attentive towards the signaller, the response elicited, and the subsequent behaviours displayed by the signaller in the absence of an immediate response. Using the intentionality criteria established by great ape researchers, we have developed a repertoire of functional gestures used by bonnet macaques, in the contexts of affiliation, aggression and play. We also evaluated the differences between mean repertoire sizes and frequencies of displayed gestures across individuals of different sexes and age classes in four study troops.

## Methods

### Study area

We conducted this study from February 2013 to July 2014 in the Bandipur National Park (11.66°N, 76.63°E) in the southern state of Karnataka, India (Fig. 1). Spanning over c. 874 km^2^, with an elevation ranging from 680 m to 1,454 m ASL, this Park experiences a typical tropical climate, prevailing across the region. The Park falls within the Nilgiri Biosphere Reserve, at the junction of the Deccan Plateau and the Western Ghats, and is characterized mostly by dry deciduous forests, interspersed with moist deciduous forest patches and dry scrub. The annual rainfall cycle allows for the demarcation of a dry and a wet season, from December to May and June to November respectively. The average rainfall in the area falls in the range of 141.44 ± 19 mm during the period from June to September (Chatterjee 2013). Bandipur is host to a rich ensemble of flora and fauna, including diverse trees, insects, amphibians, reptiles, and birds, as well as small and large mammalian species (Chatterjee 2013). The most common primate species in this area is the bonnet macaque, the study species described below. The troops that we studied belong to a free-ranging population, though leading a somewhat provisioned life along a highway that runs through the Park.

**Figure 1.**
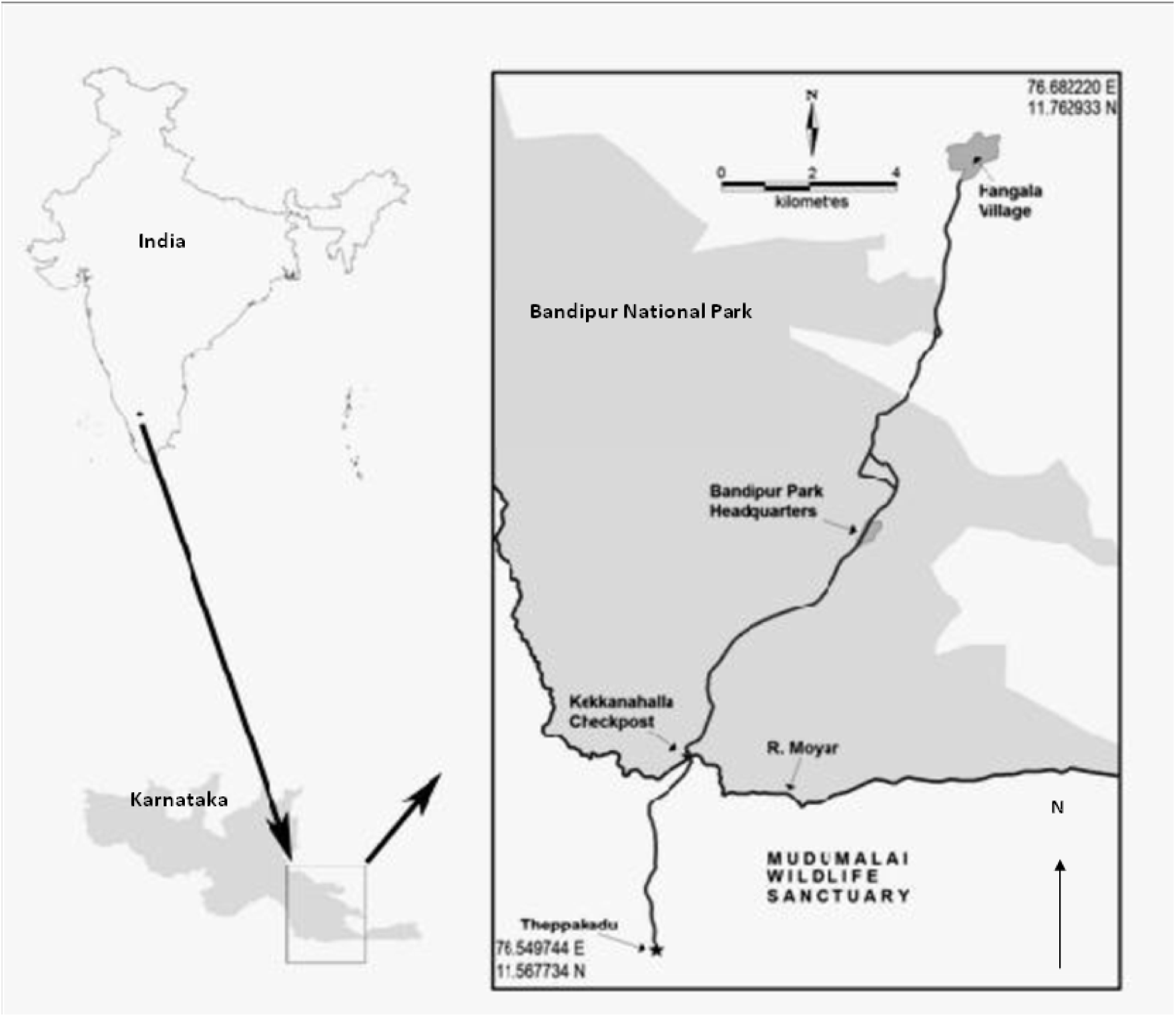
Location of the study site, the Bandipur National Park in Karnataka state, southern India (adapted from Chatterjee 2012)

### Study species

The bonnet macaque is a cercopithecine primate, endemic to peninsular India and extensively distributed across a wide range of habitats, possibly due to its exceptional ecological flexibility and behavioural lability, as has been documented in earlier studies (Sinha 2001; Sinha et al. 2005; Ram, Venkatachalam, and Sinha 2003). An elaborate behavioural repertoire and complex social interactions also characterize the species (Sinha 2001). The phenotypic plasticity displayed by this species, including the presence of behavioural traditions (Sinha 2005; Sinha et al. 2005), especially in the context of communication (Gupta, Kasturirangan, and Sinha 2015), makes this an even more suitable model system to investigate flexible systems such as gestures.

In bonnet macaque societies, females usually live in their natal troops throughout their life while males often emigrate, either transiently or permanently, to join other troops (Sinha 2001). Females form strong, linear, transitive dominance hierarchies with younger daughters of individual adult females occupying ranks just below those of their mothers. The dominance status of males is more fluid and determined mostly by aggressive interactions as well as the formation of successful coalitions (Sinha 2001).

SG identified four study troops in and around the Bandipur National Park, three of which had a species-typical multimale-multifemale social organization while one of them was a unimale-multifemale troop, an unusual, recently characterized form of social organization shown typically by this particular population (Dutta-Roy and Sinha 2001; Sinha 2001; Sinha et al. 2005; Sinha, Mukhopadhyay, and Roy 2003). The study troops comprised a total of 29 adult females (>4 years of age), 23 adult males (>4 years of age), 31 juveniles (2–4 years of age) and 26 infants (0–2 years of age) at the beginning of the study (Table 1). Infants are entirely dependent on their mothers, especially during foraging and travelling. From a year after birth until their testicles have descended (for males) or they start cycling (for females) is considered the juvenile stage, after which individuals are considered adults.

**Table 1.** Composition of the four study troops and their observation hours in the Bandipur National Park

| **Troop** | **Social organisation** | **Adult females** | **Adult males** | **Juveniles** | **Infants** |
| --- | --- | --- | --- | --- | --- |
| TT1 | Multimale-Multifemale | 7 | 7 | 9 | 6 |
| TT2 | Unimale-Multifemale | 6 | 1 | 4 | 4 |
| HN2 | Multimale-Multifemale | 8 | 5 | 5 | 9 |
| C3 | Multimale-Multifemale | 8 | 10 | 13 | 7 |
| Observation hours: |  |  | 85.75 | 100.00 | 87.50 |
| Total | | 131.75 | $3.73 \pm 0.53$ | $3.23 \pm 0.37$ | $2.88 \pm 0.44$ |
| Mean ± SE | | $4.54 \pm 0.60$ | 1.50 – 11.25 | 0.75 – 9.50 | 0.50 – 8.50 |
| Range |  | 1.50 – 11.00 |  |  |  |

### Data coding

Since 2000, AS has observed and manually documented the behaviours displayed by this population of bonnet macaques and had prepared an ethogram (Supplementary Material 1) based on the objective descriptions of behavioural events and states observed in this species (Sinha et al. 2005; Sinha and Mukhopadhyay 2013; Sinha 2001, 2003, 2005, 2014; Ram, Venkatachalam, and Sinha 2003; Dutta-Roy and Sinha 2001; Chatterjee 2013). AS trained SG in the identification of all behaviours and in the matching of the observed behaviours with those in the ethogram. SG conducted the systematic observations that manually documented gestures in this macaque population, coding of the data and all analyses. SG also collected *adlib* video recordings at the initial stage of the study. These recorded videos were scanned by AS in order to ascertain the proper identification of all the behaviours observed by SG. In order to assess inter-observer reliability, 20% (n = 36) of the recorded *adlib* videos were coded by a second researcher, who was not a part of this particular study, but has been exposed to macaque and ape behaviour studies as part of her own research. The second coder was provided with a list of potential gestures and their descriptions (Table 2). A gesture was coded only if (i) the signaller and the recipient were looking at each other, (ii) gesturing was stopped after the recipient responded, and (iii) the signaller persisted gesturing in absence of a response from the recipient. Gestures which complied with all these characteristics were then tested for reliability. The randomly chosen videos covered 81% of the reported gestures and the Cohen’s Kappa value for all the gestures ranged from 0.80 – 1.00 (except the gesture Touch, for which the reliability score was 0.56).

**Table 2.** Name, description, context and modality of the gestures displayed by bonnet macaques

| Code | Gesture | Description | Context | Modality |
| --- | --- | --- | --- | --- |
| LS | Lip-Smacking | Frequent opening and closing of lips in contact with tongue and teeth, resulting in lapping-like movement, directed toward the recipient | Affiliation | Visual |
| SG | Soliciting<br>Allogrooming | Displaying position change, head/body extension, showing rear or holding body part towards the groomer during bouts of allogrooming ((Gupta and Sinha 2016)) | Affiliation | Visual |
| BG | Biting Gently | Holding the recipient and biting lightly on its body, not closing the mouth completely | Affiliation | Tactile |
| CB | Pulling Close to Body | Holding the recipient and pulling towards the sender | Affiliation | Tactile |
| HO | Holding Gently | Holding a recipient's body part and sitting together without any other interaction | Affiliation | Tactile |
| HS | Hugging with Lip-Smacking | Encircling and holding the recipient's body with both hands along with simultaneous lip-smacking | Affiliation | Tactile |
| HU | Hugging | Encircling and holding the recipient's body with both hands | Affiliation | Tactile |
| MB | Mouth-to-Body<br>Touching | Repeatedly touching the recipient's body with the mouth | Affiliation | Tactile |
| MT | Mouth-to-Mouth<br>Touching | Coming in front of the recipient and touching the recipient's mouth with the mouth, usually once | Affiliation | Tactile |
| NZ | Nuzzling | Rubbing the face onto the recipient's body | Affiliation | Tactile |
| PA | Patting | Encircling the recipient with one hand and repeatedly patting the recipient's back/shoulder/arm | Affiliation | Tactile |
| TO | Touching | Lightly touching the recipient's body with a swift movement of the hand | Affiliation | Tactile |
| HJ | Head-Jerking | Repeated swift movements of the head, usually vertically, while looking directly towards the recipient | Agonistic | Visual |
| LU | Lunging | Sudden jump toward the recipient, followed by a swift return to the original position | Agonistic | Visual |
| OT | Open-Mouth | Opening the mouth in rounded form | Agonistic | Visual |
|  | Threatening | along with slight forward head movements<br>and followed by a rapid closing of the<br>mouth, while looking directly at the<br>recipient |  |  |
| BH | Biting Hard | Holding the recipient tightly and biting<br>with the mouth fully closed and with the<br>teeth often embedded in a recipient's body<br>part | Agonistic | Tactile |
| EF | Eye Flashing | Sudden widening of the eyes with raising<br>of the eyebrows, usually displayed<br>repeatedly, while looking directly at the<br>recipient | Agonistic | Tactile |
| HD | Holding Down<br>Roughly | Holding the recipient tightly and pushing<br>her/him down towards the ground<br>without any release | Agonistic | Tactile |
| SL | Slapping | Sudden quick movement of the palm,<br>making forcible contact with the<br>recipient's cheek/body/head | Agonistic | Tactile |
| PL | Lunging in Play | Sudden jump towards the recipient and<br>then approaching her/him, while playing | Play | Visual |
| PO | Open-Mouth | Opening the mouth in a rounded form | Play | Visual |
|  | Threatening in Play | and retaining this display, while looking at the recipient, during play |  |  |
| SJ | Spot-Jumping | Sudden, repeated jumping on the same spot, while looking at the recipient | Play | Visual |
| PB | Biting in Play | Holding the recipient and biting lightly without the mouth fully closing and often repeating this behaviour, while playing | Play | Tactile |
| PS | Slapping in Play | Slow movement of the palm, lightly brushing the recipient's cheek/head/hand/other part of the body | Play | Tactile |

### Data collection

SG and AS identified all the individuals in the study troops and categorized them into different age classes based on their body size and visually determined age-typical morphological characteristics. Following habituation and successful identification of all the individuals of a troop, SG followed a standard 15-minute focal animal sampling of all troop members without replacement (Altmann 1974). Focal sampling was carried out from 09:00 to 17:30, six days a week. If the focal animal could not be followed for more than 5 min during a sampling session, the collected data were discarded. SG recorded all individual and social behavioural events displayed by the focal subject during each session. She also monitored the contexts of the responses elicited from the recipients during all instances of behavioural signals and gestures. The context of the significantly highest proportion of the responses was considered to be the appropriate context of the displayed gestures and consequently, their function. The data consists of 392.5 hours of observation on 109 individuals including adults, juveniles and infants. The detailed description of each study troop is given in Table 1.

### Data analysis

We recorded 2,464 events of potential gesture use, when a signaller displayed a discrete signal toward a target recipient while both were looking at one another. These behavioural events were manually recorded along with the response evoked in each instance and its context. The proportions of response types displayed were compared with the G-test of independence (Sokal and Rohlf 2012) in order to establish the function/s of each initiated gesture.

Individual repertoire size for the study troop members were estimated by including all the gesture types that occurred in the contexts of affiliation, agonism and play. The frequency of gestures displayed by each individual was calculated as follows:

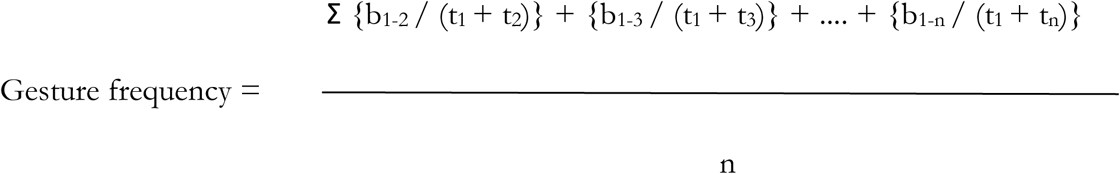

where,

b1-2 = number of gestures displayed by Individual 1 towards Individual 2
t1 = total duration of observation on Individual 1 (h)
t2 = total duration of observation on Individual 2 (h)
n = total number of individuals that received gestures from Individual 1

To test the relationship of gesture repertoire size and frequency of gesture use with age, sex and troops of the individuals, we constructed multiple statistical models. Thus, our response variables were affiliative repertoire size, agonistic repertoire size, play repertoire size, frequency of affiliative gestures, frequency of agonistic gestures and frequency of play gestures. The explanatory variables were age and sex of the individuals. Since gesture repertoire size is a count variable, we employed Laplace approximation method of generalised linear mixed models (GLMM; Poisson distribution and Log-link function) (St-Pierre, Shikon, and Schneider 2018; Bolker et al. 2009) to evaluate the differences between mean repertoire sizes in the different sexes and age classes in the four study troops. Individual identities were used as a random effect in these models. We employed linear mixed model to our frequency of gestures data. The analyses were performed within R (version 3.5.1) platform using packages MASS, nlme and lme4 (RCoreTeam 2013).

The first four criteria for a behavioural signal to be considered intentional, outlined above, were ensured in our analysis by selecting only those interactions where the signaller and the receiver were visually oriented towards one another. Thus, visual orientation was used as an initial marker for the use of intentional signaling. Additionally, we examined the goal-directedness of potential gestures displayed by the subject bonnet macaques as another marker to establish the intentional use of the observed signals. In order to achieve this, we monitored the context of the responses elicited after the display of a potential gesture. If the elicited response appeared to be appropriate for the emitted signal and did not lead to a repeated display of the same gesture or that of another signal by the signaller, we assumed that the communicative goal had been achieved (Tomasello et al. 1994). The context (affiliation/aggression/play) of the elicited response, which was displayed at the relatively highest frequency, was noted for all such interactions in order to discern the function of the particular signal. It should be noted, however, that upon sending a signal, which allowed the signaller to establish a context of communication, there could either be a response in the same context or in a different one. For example, if a signaller sent out a signal of affiliation, this could be reciprocated either by other affiliative responses (same-context responses) or by agonistic/play/subordination/sexual signals (other-context responses). A same-context response indicated that the initiated behaviour was displayed to a particular receiver with an expected outcome favourable to the signaller and that the motivation behind the displayed behaviour was considered to have been recognized. Instances that elicited responses unexpected for that particular context, as, for example, an agonistic response to an initiated affiliative behaviour, was considered to represent decisions perhaps justifiable in other situations but inappropriate in the prevailing context.

Moreover, in order to investigate the intentional use of signals used by bonnet macaques, we evaluated the situations where there was an absence of a response or an inappropriate one was displayed. These were events in which, despite the signaller and receiver looking at one another, the signaller waited for a response (typically more than 10 sec) and subsequently produced either the same behaviour persistently or other related behaviours until an appropriate response was elicited. This indicated that the signaller had an intended goal to achieve when the signal was originally sent, through which it attempted to alter the receiver’s behaviour; the failure of eliciting an appropriate response, however, motivated the signaller to further pursue this communication until the apparent goal was achieved. We followed the key criteria, established for detecting markers of intentional use of gestures by apes, in order to pursue this preliminary investigation of intentional gesture use by bonnet macaques. We monitored the appropriate responses followed by cessation of gesturing by the signallers as well as response waiting and the persistent display of gestures to establish intentionality. Other criteria of intentional use of the gestures, such as the appropriate targeting of audience and choice of modality, have not been addressed in this study and we discuss this limitation of the study below.

It must be noted in this context that we have already documented and reported intentional gestures in the context of allogrooming displayed by wild bonnet macaques towards conspecific individuals from our long-term observations of these particular study groups in the Bandipur National Park (Gupta and Sinha 2016). The current analysis is also a part of the same study. This contributes to the idea that intentional use of innate signals may be evolutionary older than earlier conceived, as has also been noticed in case of ravens (Pika and Bugnyar 2011) and fish (Vail, Manica, and Bshary 2013). Moreover, other species of macaques have been reported recently to use intentional gestures towards their human caregivers in captivity, elucidating their inherent capacity of intentional communication (Meunier, Fizet, and Vauclair 2013; Canteloup, Bovet, and Meunier 2015). We thus considered it a worthwhile exercise to investigate similar capacities but in the natural communication of individuals of a simian species in the wild.

This study involved only non-invasive, observational methods, performed on free-ranging macaques in accordance with all applicable international and national ethical guidelines and with those specified by the National Institute of Advanced Studies to which the authors are affiliated. This research thus adheres to the principles for ethical treatment of primates, as prescribed by the American Society of Primatologists, USA.

## Results

### Gestural repertoire of bonnet macaques

Following our operational definition of “gesture”—*a discrete, mechanically ineffective movement of body parts deployed intentionally with the aim of eliciting a behavioural response from the recipient*— we identified 24 gestures, recorded in 2,464 events of potential gesture use (both tactile and visual) deployed by the study population of bonnet macaques (Table 2) in different contexts, including those of affiliation, agonism and play. Of these, 12 gestures were observed in the context of affiliation, seven in agonism and five in play.

An additional eight gestures were identified in the study troops but of which we were unable to identify the function/s and context of use, as discussed below. We have, therefore, considered a total of 32 gestures to be displayed by the study bonnet macaque individuals. The repertoire size and frequencies of gesture use in contexts of affiliation, aggression and play by individuals of different age classes have been presented in Table 3.

**Table 3.** Total and contextual gestural repertoire size and frequency of gesture use displayed by individual bonnet macaques of different age classes

| **Repertoire** | **Adult**<br>**n = 52** | **Juvenile**<br>**n = 31** | **Infant**<br>**n = 26** |
| --- | --- | --- | --- |
| **Affiliative** |  |  |  |
| Mean ± SE | 11.58 ± 0.63 | 10.00 ± 0.83 | 9.27 ± 0.84 |
| Range | 7 – 19 | 1 – 18 | 3 – 17 |
| **Agonistic** |  |  |  |
| Mean ± SE | 4.87 ± 0.40 | 2.97 ± 0.44 | 1.43 ± 0.32 |
| Range | 0 – 12 | 0 – 9 | 0 – 5 |
| **Play** |  |  |  |
| Mean ± SE | 1.08 ± 0.30 | 5.16 ± 0.49 | 4.65 ± 0.32 |
| Range | 0 – 10 | 0 – 11 | 2 – 8 |
| **Total** |  |  |  |
| Mean ± SE | 17.12 ± 0.96 | 18.13 ± 1.37 | 15.19 ± 1.29 |
| Range | 6 – 35 | 3 – 36 | 4 – 28 |
| **Frequency of gestures (act/h)** |  |  |  |
| **Affiliation** |  |  |  |
| Mean ± SE | 0.77 ± 0.06 | 0.65 ± 0.05 | 0.64 ± 0.06 |
| Range | 0.26 – 2.08 | 0.19 – 1.49 | 0.32 – 1.31 |
| **Agonism** |  |  |  |
| Mean ± SE | 0.30 ± 0.03 | 0.23 ± 0.05 | 0.11 ± 0.03 |
| Range | 0 – 0.83 | 0 – 1.07 | 0 – 0.57 |
| **Play** |  |  |  |
| Mean ± SE | 0.18 ± 0.05 | 0.78 ± 0.11 | 0.91 ± 0.13 |
| Range | 0 – 1.81 | 0 – 2.52 | 0.22 – 2.45 |
| **Total** |  |  |  |
| Mean ± SE | 1.25 ± 0.09 | 1.73 ± 0.12 | 1.66 ± 0.17 |
| Range | 0.44 – 3.48 | 0.74 – 3.64 | 0.65 – 3.60 |

### Effect of age on gesture repertoire size

The affiliative and agonistic repertoire sizes of infant bonnet macaques were significantly smaller. In juveniles, the same trend holds true, only in case of the agonistic repertoire size. Play repertoire size, on the other hand, was significantly larger in infants and juveniles both (Fig. 2; Table 4).

**Figure 2.**
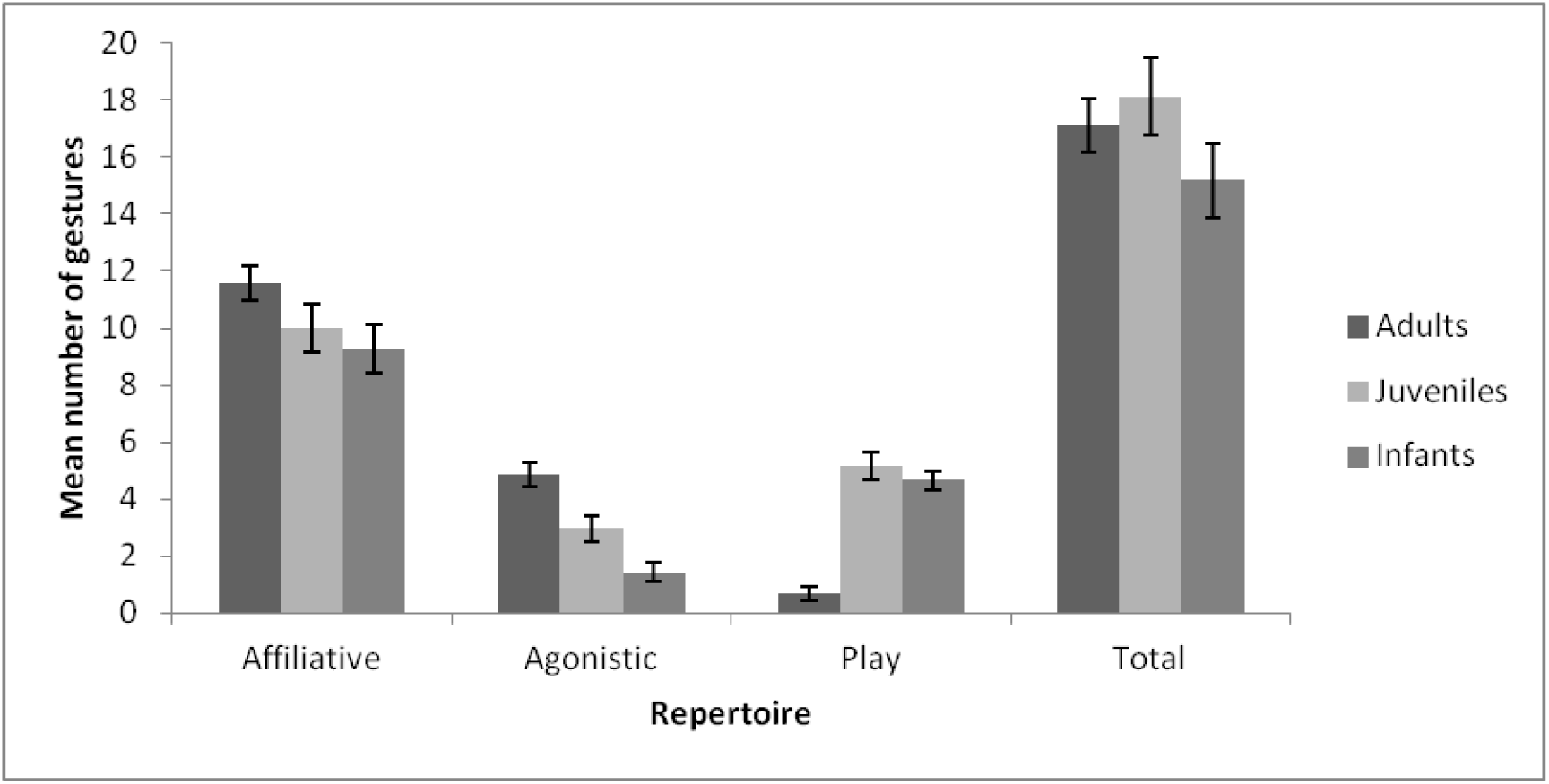
Total and contextual gestural repertoire size of adult, juvenile and infant bonnet macaques during the study period

**Table 4.** Effect of age, sex and troop on gesture repertoire size and frequency of gesture use in bonnet macaques.

|  |  |  |  |  |
| --- | --- | --- | --- | --- |
| **Affiliative repertoire** |  |  |  |  |
| **Fixed effects (Generalised Linear Mixed Model)** |  |  |  |  |
|  | **Estimate** | **Standard error** | **Z value** | **p-value** |
| **Intercept** | 2.31706 | 0.07823 | 29.617 | **<0.0001** |
| **Infant** | -0.22690 | 0.08474 | 2.678 | **0.007** |
| **Juvenile** | -0.14903 | 0.07842 | -1.900 | 0.0573 |
| **Male** | -0.37819 | 0.07004 | -5.400 | **<0.0001** |
| **HN2** | 0.39247 | 0.09015 | 4.354 | **<0.0001** |
| **TT1** | 0.47679 | 0.08620 | 5.531 | **<0.0001** |
| **TT2** | 0.18000 | 0.11051 | 1.629 | 0.10337 |
| **Agonistic repertoire** |  |  |  |  |
| **Fixed effects (Generalised Linear Mixed Model)** |  |  |  |  |
|  | **Estimate** | **Standard error** | **Z value** | **p-value** |
| **Intercept** | 1.2854 | 0.1543 | 8.328 | **<0.001** |
| **Infant** | -1.4155 | 0.2134 | -6.633 | **<0.001** |
| **Juvenile** | -0.4721 | 0.1563 | -3.021 | **0.0025** |
| **Male** | -0.3447 | 0.1420 | -2.427 | **0.015** |
| **HN2** | 0.6867 | 0.1787 | 3.843 | **0.0001** |
| **TT1** | 0.5095 | 0.1760 | 2.894 | **0.0038** |
| **TT2** | 0.1656 | 0.2280 | 0.726 | 0.4677 |
| **Play repertoire** |  |  |  |  |
| **Fixed effects (Generalised Linear Mixed Model)** |  |  |  |  |
|  | **Estimate** | **Standard error** | **Z value** | **p-value** |
| **Intercept** | -0.7978 | 0.2630 | -3.034 | **0.002** |
| **Infant** | 1.5992 | 0.2298 | 6.958 | **<0.001** |
| **Juvenile** | 1.7871 | 0.2216 | 8.066 | **<0.001** |
| **Male** | 0.5182 | 0.1656 | 3.130 | **0.0018** |
| **HN2** | 0.5438 | 0.2176 | 2.499 | **0.012** |
| **TT1** | 0.6485 | 0.2030 | 3.195 | **0.00140** |
| **TT2** | 0.2629 | 0.2770 | 0.949 | 0.34244 |
| **Frequency of affiliative gestures** |  |  |  |  |
| **Fixed effects (Linear Mixed Model)** |  |  |  |  |
|  | **Value** | **Standard Error** | **t-value** | **p-value** |
| **Intercept** | 1.0189087 | 0.07032126 | 14.489341 | **<0.0001** |
| **Infant** | -0.0894628 | 0.07712912 | -1.159910 | 0.2488 |
| **Juvenile** | -0.1535845 | 0.07296086 | -2.105026 | **0.037** |
| **HN2** | -0.3304426 | 0.08185659 | -4.036848 | **0.0001** |
| **TT2** | 0.0314002 | 0.09984466 | 0.314490 | 0.7538 |
| **Male** | -0.2087975 | 0.06360935 | -3.282497 | NaN |
| **Frequency of agonistic gestures** |  |  |  |  |
| **Fixed effects (Linear Mixed Model)** |  |  |  |  |
|  | **Value** | **Standard Error** | **t-value** | **p-value** |
| **Intercept** | **0.4576554** | **0.03975106** | **11.513038** | **<0.0001** |
| **Infant** | -0.1690660 | 0.04366983 | -3.871459 | **0.0002** |
| Juvenile | -0.0186475 | 0.04169954 | -0.447187 | 0.6557 |
| HN2 | -0.1936866 | 0.04576048 | -4.232617 | **0.0001** |
| TT2 | -0.1317298 | 0.05634675 | -2.337842 | 0.0213 |
| Male | **-0.1003500** | **0.03364093** | **-2.982973** | **NaN** |
| Frequency of play gestures |  |  |  |  |
| Fixed effects (Linear Mixed Model) |  |  |  |  |
|  | Value | Standard Error | t-value | p-value |
| Intercept | 0.1147863 | 0.1146037 | 1.001594 | 0.3189 |
| Infant | 0.6935445 | 0.1256985 | 5.517523 | **<0.0001** |
| Juvenile | 0.6033625 | 0.1189054 | 5.074305 | **<0.0001** |
| HN2 | -0.1296119 | 0.1334030 | -0.971582 | 0.3335 |
| TT2 | 0.3446783 | 0.1627184 | 2.118250 | 0.036 |
| Male | 0.2204098 | 0.1036652 | 2.126171 | NaN |
| Effect of sex on frequency of affiliative gestures |  |  |  |  |
| Linear Regression |  |  |  |  |
|  | Estimate | Standard | t-value | p-value |
|  |  | **Error** |  |  |
| **Male** | -0.20880 | 0.06361 | -3.282 | **0.0014** |
| **Effect of sex on frequency of agonistic gestures** |  |  |  |  |
| **Male** | -0.11371 | 0.03616 | -3.145 | **0.0022** |
| **Effect of sex on frequency of playgestures** |  |  |  |  |
| **Male** | 0.2204 | 0.1037 | 2.126 | **0.0359** |

### Effect of age on frequency of gesture use

The frequency of use affiliative gestures and that of agonistic gestures were significantly lower in the juveniles and infant individuals, respectively. Frequency of play gestures were, however, significantly higher in both of these two age classes. The affiliative repertoire sizes were significantly more in troops TT1 and HN2, while agonistic and play repertoire sizes were significantly more in TT1 (Fig. 3; Table 4).

**Figure 3.**
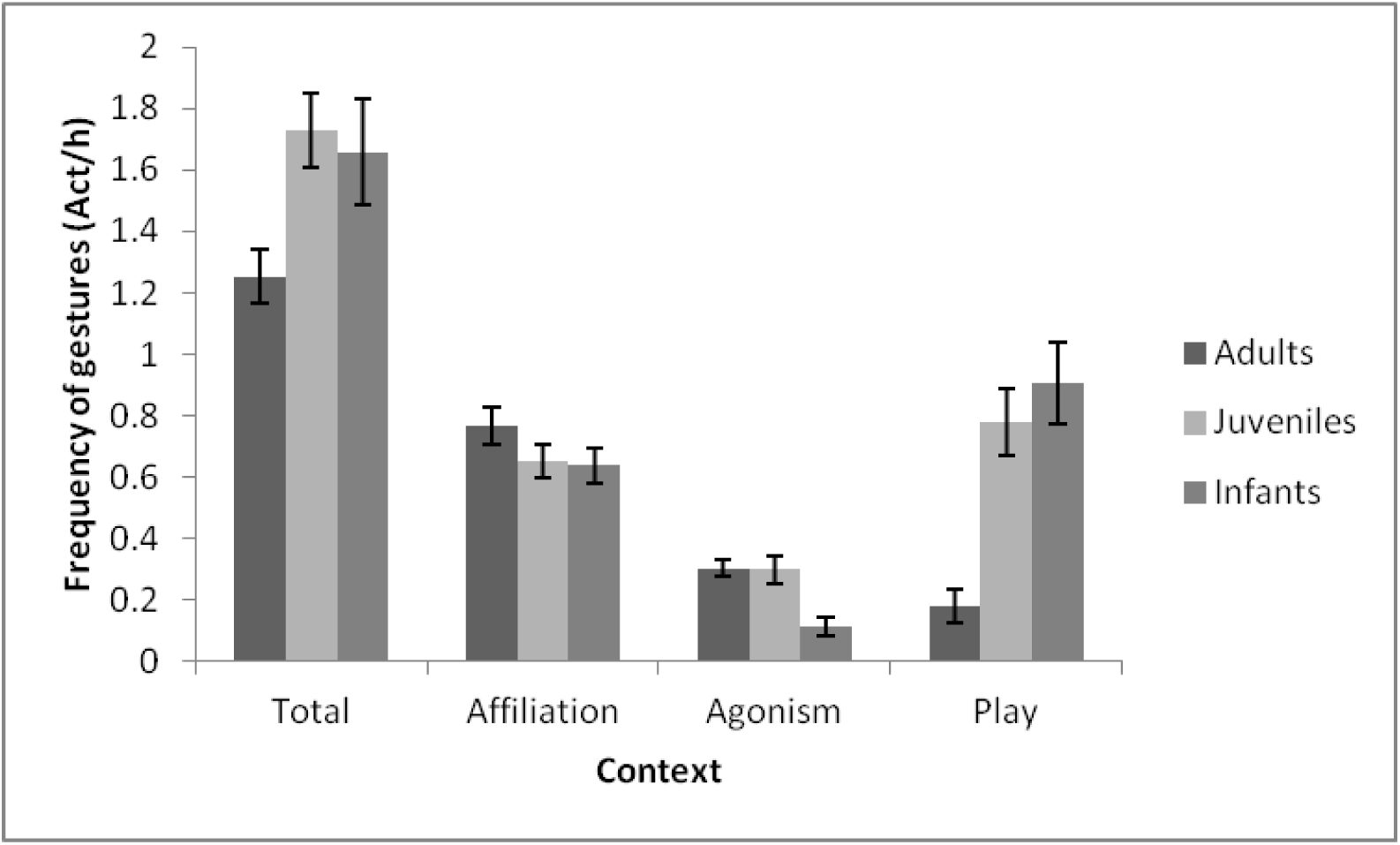
Mean frequencies of gestures displayed by adults, juveniles and infants of the four study troops in different contexts. The error bars represent standard error of the mean

### Effect of sex on gesture repertoire size and frequency of gesture use

The male individuals, irrespective of age class, showed a significantly smaller affiliative and agonistic repertoire size, while a significantly larger play repertoire size. In case of frequency of gestures displayed, the GLMM could not estimate the effect of sex on the frequency of gesture use in the different contexts. We thus used linear regression model to estimate only this effect. The males displayed significantly lower frequencies of affiliative and agonistic gestures, but higher frequencies of play gestures. Frequency of affiliative gestures were lower in HN2, those of agonistic gestures were lower in HN2 and TT2, while play gesture frequencies were higher in TT2.

### Elicited responses and contexts of the gestures

Of the 32 gestures observed, three signals (Biting Hard, Spot-Jumping and Hugging with Lip-Smacking) elicited responses in the same context almost exclusively (Fig. 4). The first two elicited agonistic and play responses respectively in all instances of their occurrence. Hugging with Lip-Smacking evoked affiliative responses on 94.6% of instances while the remaining elicited either Moving Away or Avoiding, both considered neutral responses; there were no other-context responses to this signal. The percentage of affiliative responses to Hugging with Lip-Smacking was, however, significantly higher than that of the neutral responses (G-test of independence, df = 1, G = 317.60, *p* < 0.001, n = 37; Fig. 4).

**Figure 4.**
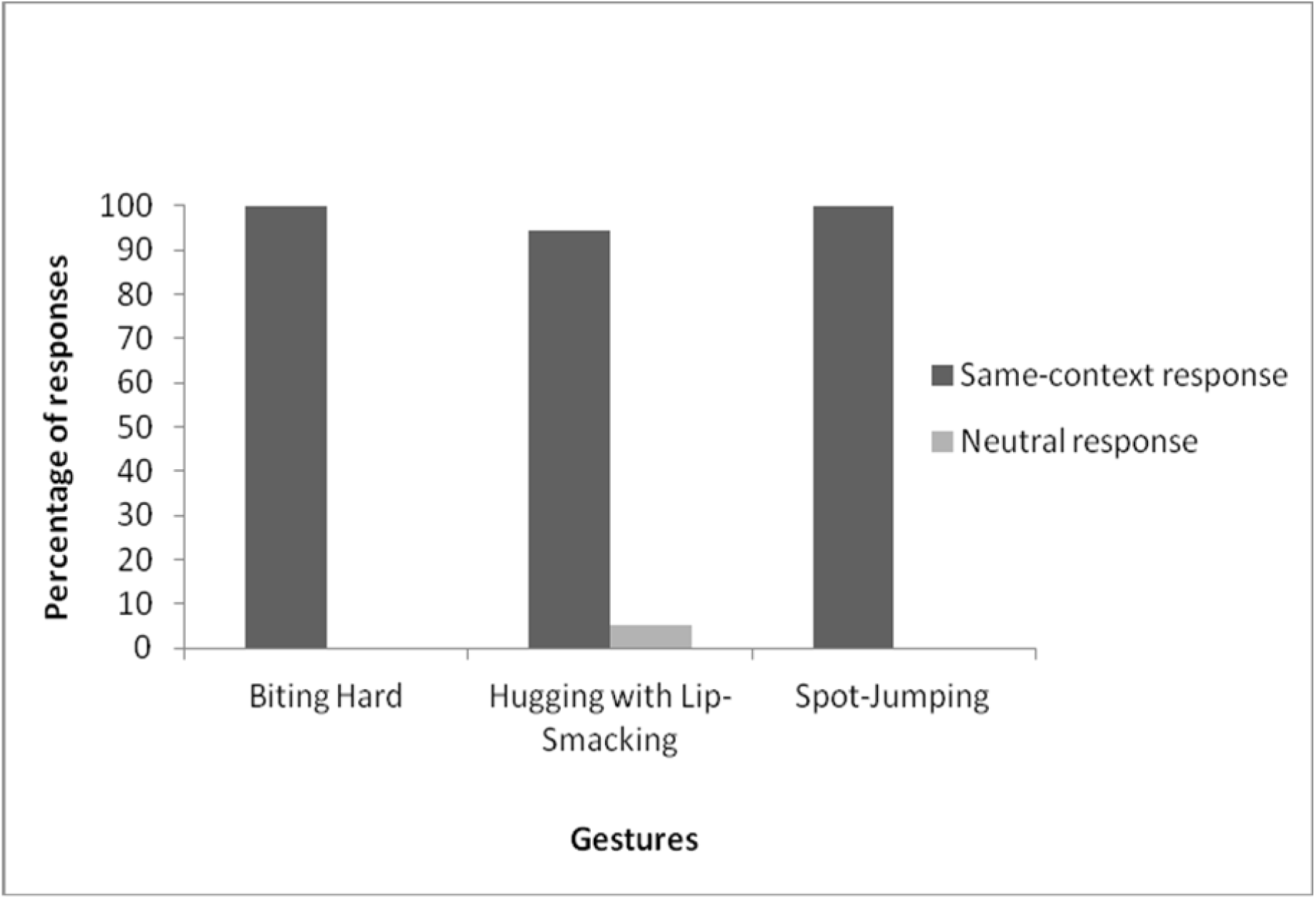
Three gestures that invariably elicited same-context responses: Biting Hard (n = 2), Hugging with Lip-Smacking (n = 47) and Spot-Jumping (n = 6)

In the case of the other 21 gestures, responses were elicited across multiple contexts and we evaluated the proportions of responses elicited by each gesture in these different contexts. We then tested, for each gesture, whether the highest proportion of responses in a particular context occurred significantly more than would be expected by chance and thus evaluated the possible function of each gesture. The various responses elicited for each gesture, their proportions and comparison for significance has been presented in Table 5. The contexts of 12 affiliative gestures, seven agonistic and five play gestures in the repertoire of bonnet macaques were thus derived from the responses elicited in the recipients (Fig. 5).

**Figure 5.**
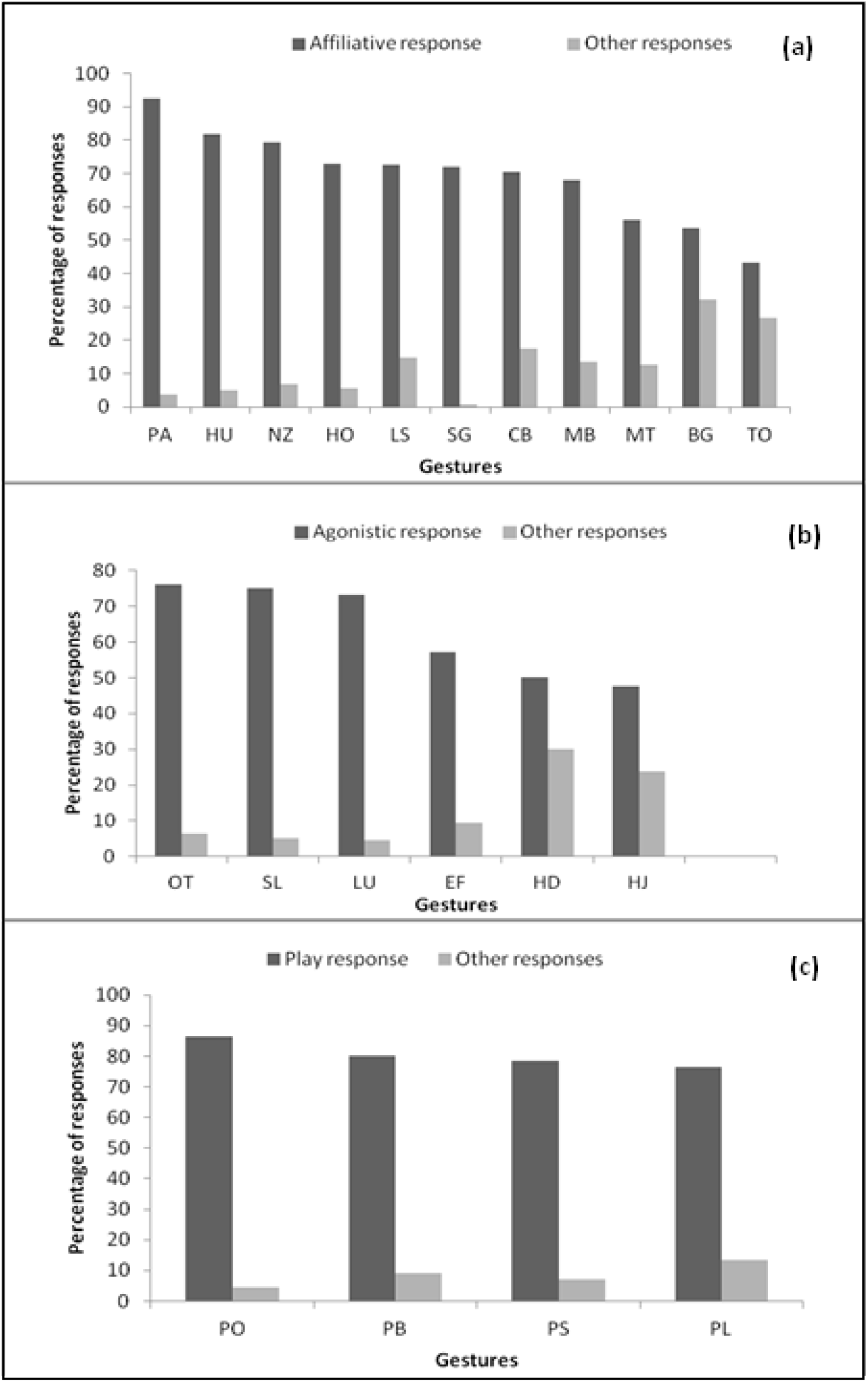
Percentage of responses in different contexts to (a) Affiliative, (b) Agonistic, and (c) Play gestures. The codes for the different gestures have been listed in Table 2.3

**Table 5.** Responses to different gestures in the contexts of affiliation, agonism and play

| **Code** | **Gesture** | **Highest**<br>**response**<br>**(%)** | **Other**<br>**responses**<br>**(%)** | **Neutral**<br>**response**<br>**(%)** | **G** | ***p*** | **n** |
| --- | --- | --- | --- | --- | --- | --- | --- |
| **Affiliative gestures** |  |  |  |  |  |  |  |
| PA | Patting | 92.59 | 3.70 | 3.70 | 674.84 | < 0.001 | 27 |
| HU | Hugging | 81.88 | 5.00 | 13.13 | 386.11 | < 0.001 | 158 |
| NZ | Nuzzling | 79.31 | 6.90 | 13.79 | 318.17 | < 0.001 | 29 |
| HO | Holding | 72.86 | 5.71 | 21.43 | 284.82 | < 0.001 | 70 |
| LS | Lip-Smacking | 72.73 | 14.69 | 12.59 | 202.65 | < 0.001 | 143 |
| SG | Soliciting<br>Allogrooming | 72.09 | 0.58 | 27.33 | 695.70 | < 0.001 | 406 |
| CB | Pulling Close<br>to Body | 70.59 | 17.65 | 11.76 | 185.47 | < 0.001 | 17 |
| MB | Mouth-to-Body<br>Touching | 68.18 | 13.64 | 18.18 | 156.84 | < 0.001 | 24 |
| MT | Mouth-to-Mouth<br>Touching | 56.25 | 12.5 | 31.25 | 104.42 | < 0.001 | 16 |
| BG | Biting Gently | 53.57 | 32.14 | 14.29 | 81.80 | < 0.001 | 28 |
| TO | Touching | 43.14 | 26.80 | 30.07 | 12.72 | < 0.001 | 161 |
| **Agonistic gestures** |  |  |  |  |  |  |  |
| OT | Open-Mouth<br>Threatening | 76.09 | 6.52 | 17.39 | 291.40 | < 0.001 | 46 |
| SL | Slapping | 75 | 5 | 20 | 319.40 | < 0.001 | 20 |
| LU | Lunging | 73.13 | 4.48 | 22.39 | 323.84 | < 0.001 | 67 |
| EF | Eye-Flashing | 57.14 | 9.52 | 33.33 | 142.79 | < 0.001 | 21 |
| HD | Holding<br>Down<br>Roughly | 50 | 30 | 20 | 42.14 | < 0.001 | 10 |
| HJ | Head-Jerking | 47.62 | 23.81 | 28.57 | 26.79 | < 0.001 | 21 |
| **Play gestures** |  |  |  |  |  |  |  |
| PO | Open-Mouth<br>Threatening in<br>Play | 86.36 | 4.55 | 9.09 | 467.77 | < 0.001 | 22 |
| PB | Biting in Play | 80.26 | 9.21 | 10.53 | 311.98 | < 0.001 | 75 |
| PS | Slapping in<br>Play | 78.57 | 7.14 | 14.29 | 306.08 | < 0.001 | 14 |
| PL | Lunging in<br>Play | 76.38 | 13.57 | 10.05 | 253.70 | < 0.001 | 199 |

Gestures were classified as affiliative, agonistic and play on the basis of the context in which they each received the highest proportion of responses. The
proportions of responses received by each gesture were compared to what could be obtained by chance alone using the G-test of independence ( $df = 1$ )

The remaining eight gestures elicited responses that did not allow us to identify particular contexts to which they belonged. The different responses, belonging to different contexts for each gesture, have been presented in Table 6. It is evident that there was not a single context, of any of these responses, which emerged to be significantly higher in occurrence than the rest. It is possible that these gestures served specific functions in different contexts and we have, thus, considered them to be multifunctional in nature.

**Table 6.** Multifunctional gestures that failed categorisation due to lack of predominant responses in a particular category

| Code | Gesture | Affiliative<br>response<br>(%) | Agonistic<br>response<br>(%) | Play<br>response<br>(%) | Dominance-<br>subordination<br>response<br>(%) | Sexual<br>response<br>(%) | Neutral<br>response<br>(%) | n |
| --- | --- | --- | --- | --- | --- | --- | --- | --- |
| MO | Mounting | 38.46 | 7.69 | 7.69 | 15.38 | 0 | 30.77 | 13 |
| PR | Presenting | 1.01 | 2.02 | 0 | 38.38 | 32.32 | 26.26 | 99 |
| PT | Pulling any<br>Part of the<br>Body | 17.48 | 30.77 | 25.17 | 1.40 | 0.7 | 24.48 | 141 |
| PU | Pushing | 14.89 | 34.04 | 17.02 | 2.13 | 0 | 31.91 | 47 |
| PN | Pinching | 0 | 44.44 | 33.33 | 0 | 0 | 22.22 | 9 |
| ST | Staring | 0 | 23.08 | 3.85 | 15.38 | 0 | 57.69 | 15 |
| RB | Raising<br>Eyebrows | 25 | 25 | 0 | 0 | 0 | 50 | 8 |
| CS | Copulatory<br>Lip-Smacking | 22.73 | 36.36 | 0 | 9.09 | 4.55 | 27.27 | 22 |

### Goal-directedness of the gestures

In 91 out of the 2,464 instances in which a signaller initiated a single gesture, the receiver did not respond despite them looking at one another continuously during the event. In the initial absence of a response, the signallers typically waited for more than 10 sec and repeatedly displayed either the same or a different gesture, targetted to the original receiver. A positive response was eventually evoked in 88 (96.7%) of these instances while on three occasions, there was ultimately no response from the receiver. Of the 88 positive responses, 13 (14.8%) and 9 (10.2%) were either in other contexts or constituted neutral responses respectively. An appropriate response in the same context of the signal was elicited in the majority of these instances (66 or 75% of 88 instances; G-test of independence, df = 1, G = 236.85, *p* < 0.001).

## Discussion

### Gestural repertoire of bonnet macaques

Our present study is a preliminary but systematic documentation of the gestural repertoire of wild bonnet macaques, following several criteria that have been put forward to define gestures and establish the underlying intentionality of their production in earlier studies on nonhuman primates. Given the preliminary nature of this investigation, it is entirely possible that the complete repertoire of gestures of bonnet macaques may not be represented in our study. Moreover, our stringent application of the criteria by which a signal could be considered a gesture may have led to the exclusion of certain gestures from our current list. Each of the gestures reported in this study, however, met the conditions of qualifying as a gesture.

There are, however, certain caveats that must be remembered in this context. First, for instance, the adjustment of the signaller to the attention states of the recipients and gaze alternation between the signaller and the goal of communication could not be directly tested in this study. Instead, we only considered those behavioural signals where the signaller and the receiver were already visually oriented towards one another while ensuring that these signals fulfilled the other necessary criteria. Our next step would be to investigate all other criteria of intentional use of gestures by bonnet macaques, which could not be appropriately addressed here. The criteria for goal-directed communication or intentional use of gestures have so far been experimentally verified for apes and monkeys in captivity (Leavens, Russell, and Hopkins 2005; Roberts et al. 2014; Cartmill and Byrne 2010; Canteloup, Bovet, and Meunier 2015, 2015) and through observations of natural communication behaviour exclusively of apes (Hobaiter and Byrne 2011; Genty et al. 2009; Byrne et al. 2017; Pika, Liebal, and Tomasello 2003, 2005; Liebal, Pika, and Tomasello 2006, 2004). The present study, therefore, becomes the primary descriptive source of gestural signals for any non-ape species under natural conditions, with the potential of facilitating future comparative studies on various aspects of gestural communication in non-ape simian species.

Our study bonnet macaques displayed a repertoire of 24 unique gestures in the contexts of affiliation, agonism and play. Moreover, eight other gestures, the contexts of which could not be determined, also featured in the gestural repertoire of this population. As clarified above, all 32 gestures were successfully tested for criteria adapted from earlier studies on human and nonhuman primate communication (Genty et al. 2009; Hobaiter and Byrne 2011a; Call and Tomasello 2007; Bates, Camaioni, and Volterra 1975; Piaget and Cook 1952; Plooij 1977; Bard 1992; Leavens and Hopkins 1998; Townsend et al. 2017). In each case, the gestures were invariably displayed by signallers while the target recipients were directly looking at them. In most instances, the receiver responded immediately to the signal but in the absence of an initial response, the signaller persistently displayed the same or a related gesture until an appropriate response was elicited and received.

The study bonnet macaques used both visual and tactile gestures. An additional category of primate auditory gestures has also been reported earlier (Tomasello et al. 1994; Pika, Liebal, and Tomasello 2003; Liebal, Pika, and Tomasello 2006; Call and Tomasello 2007; Genty et al. 2009; Hobaiter and Byrne 2011a) but this category was difficult to define in our study species, as the sound produced during the display of some of these gestures seemed to be a by-product of the actual behaviour shown and did not appear to independently function as an auditory signal. For instance, gestures such as Lip-Smacking, Hugging with Lip-Smacking, Patting, Lunging, Slapping, Lunging in Play, and Slapping in Play often produced an audible sound on contact with other individuals or with the substratum on which the behaviours were displayed see also (Hobaiter and Byrne 2011a). It is interesting to note here that the macaque hardly use manual gestures, contrasting to the apes. Walking on four limbs, almost always, could be restricting the forms of gesture production in monkeys. A gradual pattern of evolution gesture forms or manual gesturing from monkeys to apes could be potential line of investigation in future.

### Age and sex differences in repertoire size and frequency of use of gestures

The complete gestural repertoire size of adult, juvenile and infant bonnet macaques were not significantly different from each other, unlike in chimpanzees (Hobaiter and Byrne 2011; Tomasello et al. 1997; Tomasello et al. 1994), bonobos ((Pika, Liebal, and Tomasello 2005)), gorillas (Genty et al. 2009; Pika, Liebal, and Tomasello 2003), orangutans (Liebal, Pika, and Tomasello 2006), siamangs (Liebal, Pika, and Tomasello 2004) and Barbary macaques (Hesler and Fischer 2007). While the gestural repertoire of juvenile bonnet macaques included a few gesture types not displayed by the adults, their repertoire size was not significantly different. Amongst monkeys, the gestural repertoire size of Barbary macaques has been reported to be the largest for adults (Hesler and Fischer 2007).

It is noteworthy that when the complete repertoire of the study species was categorized on the basis of the different contexts of gesture production, the affiliative and agonistic gesture repertoires showed a significant increase in size with age, with adults having larger repertoires than either the juveniles or infants. In the case of the play gestural repertoire, however, repertoire size significantly decreased with age, with infants having the highest number of play gestures that gradually reduced in number in juveniles and in adults. Although the overall repertoire size increased with age, our finding that the play repertoire decreased with age is compatible with those from other species, where the overall decrease in repertoire size was also largely due to the gestures used in play. This result differs, however, from that in apes, wherein juveniles are known to have the largest repertoire amongst all age classes, resulting from a significantly varied play repertoire in the respective species (chimpanzees: (Tomasello et al. 1997; Hobaiter and Byrne 2011; Tomasello et al. 1994); bonobos: (Pika, Liebal, and Tomasello 2005); gorillas: (Genty et al. 2009; Pika, Liebal, and Tomasello 2003); orangutans: (Liebal, Pika, and Tomasello 2006).

It is important to note here that although each age class of bonnet macaques had comparable total gestural repertoire sizes, the contextual repertoire size varied with age. It is obvious that the contexts of social interactions change with progressing age and this is likely to have given rise to the differential nature of the gestural repertoires displayed by each age class. Thus, the lower number of affiliative and agonistic gestures in infant and juvenile bonnet macaques is compensated by a greater number of play gestures, an activity in which they spent most of their observed time.

Among males, affiliative and agonistic repertoire sizes were significantly smaller than females, while play repertoire was significantly larger. In the gestural communication system of other primates, there are repertoire differences among the two sexes as well as category preferences, as has been reported in several species. Specific gestures, as, for instance, in sexual contexts, are typically displayed only by male siamangs and Barbary macaques (Hesler and Fischer 2007; Liebal, Pika, and Tomasello 2004). Female orangutans also display more visual gestures than do males (Liebal, Pika, and Tomasello 2006) while male orangutans as well as siamangs produced relatively more tactile gestures (Liebal, Pika, and Tomasello 2006, 2004). Gestural studies in chimpanzees have rarely investigated such issues except for certain observations on sex differences in sexual signalling (Hobaiter and Byrne 2012) or signalling to humans under captive conditions (Hopkins and Leavens 1998). A recent study by Scott (2013) focussed systematically on sexual differences in gesture use by chimpanzees and illustrate how evolutionarily distinct pressures could have potentially shaped isolated repertoires for each sex. I made similar observations in bonnet macaques, wherein females, in general, had larger repertoires than did males, possibly contributed to by the larger inventory of affiliative gestures that they typically exhibited. This strongly supports the hypothesis put forward by Scott (2013) that social pressures may be acting differently on the two sexes, resulting in distinct communication strategies in female and male individuals. Bonnet macaques, typically forming female-bonded societies, are likely to consist of females who would display a greater diversity of affiliative gestures than would males. The social hierarchies of the study troops or the age differences among the adults in them, however, did not appear to influence the gestural repertoire size, unlike that reported in other macaques (Maestripieri 2005).

The frequency of use of all gestures taken together, by bonnet macaques, does not vary with their age class. However, it has been reported for several ape species that juveniles display higher frequencies of gestures, overall (chimpanzees: (Tomasello et al. 1997; Tomasello et al. 1994; Hobaiter and Byrne 2011); bonobos: (Pika, Liebal, and Tomasello 2005); gorillas: (Genty et al. 2009; Pika, Liebal, and Tomasello 2003); orangutans: (Liebal, Pika, and Tomasello 2006). Individual bonnet macaques of various age classes displayed a variable use of contextual gestures in affiliation, agonism and play. Affiliative gestures were significantly less frequent among juveniles, while infants showed lower levels of agonistic gestures. Infants, on the other hand, exhibited the highest frequency of play gestures, followed by juveniles, with the levels of these gestures declined significantly among the adults.

When the total frequency of gesture use was considered, infants displayed the highest frequency of play and affiliative gestures in their communication repertoire. These were accompanied by the gradual appearance of agonistic gestures at the juvenile stage while, finally, the adult repertoire predominantly consisted of affiliative and agonistic gestures. Thus, the juvenile stage appeared to be the age at which bonnet macaques used most of the gesture categories at comparable levels in their repertoire, possibly leading to the highest frequency of total gesture use amongst the three age classes studied.

Sex differences in the use of gestures were also prominent in case of adult bonnet macaques, similar to the observations made for their respective repertoire sizes as well. The frequency of use of all gestures across contexts was not significantly different amongst adult females and males; however, females displayed higher frequencies of both affiliative and agonistic gestures than did males. These high levels of gestural display were possibly compensated by the relatively high frequency of play gestures displayed by adult males, thus yielding comparable levels of total gestural frequencies by the two sexes. The use of play gestures by male bonnet macaques possibly owed much to the subadult males, who were included in this category and who often engaged in high levels of play behaviour. Sexual differences in gesture use have also been reported in other primates such as the pigtailed macaque, wherein the frequency of certain gestures, as, for instance mounting or headstands, were relatively more displayed by males than by females (Maestripieri 1996, 1997) or in chimpanzees, where males exhibited higher frequencies of consortship gestures than did females (Hobaiter and Byrne 2012). This is perhaps in concordance with the hypothesis proposed by (Scott 2013) that differential selective pressures on females and males may have resulted in the differential use of gestures by the two sexes in primates, and the bonnet macaque may be no exception.

A perspective of gradual development of the gestural repertoire and its usage is indicated by these age-specific differences in repertoire size and frequency of gesture use in bonnet macaques. The relatively lower number of contextual gestures in the infant and juvenile repertoires, especially the affiliative and agonistic gestures, probably indicates the requirement of a phase of maturation before individuals can actively produce these functional gesture forms. This phase could thus possibly be compared to the first level of cognitive developmental stage in humans, as described in (Piaget and Cook 1952). The juvenile stage possibly represents the beginning of behavioural independence, when individuals move around freely and encounter various social contexts where the newly emerging signals could be used. We believe that this stage, bonnet macaque individuals are in the process of polishing their gestural repertoire and learning the appropriate signals for each context as well as their functions, akin to what has been suggested for juvenile apes, the so-called “repertoire tuning hypothesis” discussed in (Byrne et al. 2017). This could be comparable to the crucial next Piagetian stage in humans, where an appropriate means-end use of gestures sets in to achieve particular intended communicative functions. Such goal-directed behaviour meets the crucial criteria of intentionality underlying gestural signals, a traditional goal of studies in primate communication see (Call and Tomasello 2007). We also noticed that adult individuals did not to display certain gestures, for instance, play gestures, though it was evident that they were quite capable of producing them, as expressed during occasional play sessions with juveniles. It, therefore, seems apparent that an innate set of signals, some of which first appear in infancy, go through various stages of physio-psychological development, giving rise to age-specific repertoires and the usage of certain kinds of context-appropriate gestures during different developmental stages in the natural communication system of the species.

Differences in gesture repertoire size and frequency of use was noticed in the various troops, the reasons of which could not be ascertained properly. It is, however, worth mentioning that the local habitat conditions were variable across troops, some with more vegetation, some with large open spaces and some in closer proximity to human settlements than others. It is perhaps indicative that local ecological conditions influence the communication behaviour at the troop level in bonnet macaques.

It must be mentioned here that during this study, we also observed other gesture-like behaviours, which occurred in sequences and evoked specific responses but it was not possible to determine the correspondence between these responses and the constituent behaviours within each sequence. These behaviours, therefore, could not be considered as single gestures in accordance with the strict definitional criteria that we have followed in this paper. An understanding of these gesture sequences, nevertheless, will be crucial to understanding the complete gestural repertoire of the species. Such sequences have already been recorded in apes (Hobaiter and Byrne 2011a, 2011b; Genty et al. 2009; Genty and Byrne 2010; Liebal, Call, and Tomasello 2004; Call and Tomasello 2007; Tomasello et al. 1994) and it is imperative that they be analysed in bonnet macaques as well in the future.

### Elicited responses and contexts of the gestures

The three gestures—Biting Hard, Spot-Jumping and Hugging with Lip-Smacking—which always received responses in the same relevant context, appeared to be intentionally produced in agonistic, play and affiliative contexts respectively. The contextuality of the other 21 gestures, the functions of which could be clearly defined, were similarly established but only by the significantly higher proportions of responses elicited in those particular contexts. The predominant response type indicated particular communicative functions of these gestures, similar to those observed in orangutan gestures, where typical gestures were stably associated with their contexts, representing the semantics of the signal (Cartmill and Byrne 2007). The responses evoked in the other minor contexts could possibly be attributed to a host of putative factors, including the motivational states of particular recipients, interactions preceding the signalling interactions and previous to our observations, developmental factors or even the social history of the participants of the communicative events.

The functions of the additional eight gestures remain undefined owing to the absence of predominant response types. These gestures were indeed targeted to particular recipients, visually oriented towards the signaller and used intentionally, as evident from the persistent gesturing displayed by the signaller. Mounting, for instance, elicited the highest percentage of affiliative responses but this was not significantly different from those elicited in other contexts. It is possible that these gestures with undecided functions were more often used in association with other gestures, a hypothesis not tested in the present analysis. Additionally, these gestures could potentially be of flexible nature, thus making them ubiquitously applicable across contexts, as has been observed in apes as well (Tomasello and Call 2007; Tomasello et al. 1994; Tomasello et al. 1997; Genty et al. 2009; Hobaiter and Byrne 2011a). It is also possible that these gestures could even serve innovative, independent functions in different troops, practices that may then be transmitted culturally, as has been demonstrated for a particular signal, branch-shaking, in another population of bonnet macaques (Sinha 2005). It has also been provocatively suggested that certain behaviours may be in the process of becoming established as ritualized signals in particular contexts (Daanje 1951; Tinbergen 1952; Morris 1957; Lorenz 1966; Zahavi 1980) and the function of such ritualization could become less ambiguous over time in the history of a species (Cullen 1966). Is it then possible that these apparently multifunctional gestures have not yet assumed definitive roles but would become ritualized between signallers and receivers by mutual understanding over evolutionary timescales in order to serve a certain function in a particular context?

### Goal-directedness of the gestures

That bonnet macaques intentionally used gestures was evident when they persistently displayed the same gesture or other gestures upon failure to elicit a response from the recipient in the first attempt. A significantly high proportion of such persistent gesturing ultimately evoked appropriate responses from the recipient, indicating the goal-orientedness of the signaller’s behaviour in the first place see (Tomasello and Call 2007). It is worth mentioning here that individuals from this same population of bonnet macaques are capable of intentional signalling through gestures and vocalisations in a unique context of requesting food from humans (Deshpande, Gupta, and Sinha 2018). Occasionally, when neutral responses were elicited, however, it was not possible to understand the intentionality behind those particular gesturing events. In the present study, we have explored only the signaller’s wait for a response and her persistence behaviour as markers of intentionality underlying bonnet macaque gestures. There, thus, remains an extensive scope of investigating other criteria of intentionality, such as gaze alternation and adjustments made in response to the audience’s attention state, in order to holistically understand intentional use of gestural communication in this species. Future studies using video recording will offer a more fine-grained analysis of macaque gestures, allowing us to include more gestures within our analysis of intentional usage. It must be noted here, however, that the idea of intentionality, especially in species usually associated with lesser cognitive complexity, has always been questioned. Following generalized criteria of intentionality, nevertheless, we know that ravens (Pika and Bugnyar 2011) and even fish (Vail, Manica, and Bshary 2013) are capable of such intentional communication. It may thus be time for a change of perspective: in accepting that the capacity of intentional communication is evolutionarily older than we had previously thought and that it may not have been extremely complex to begin with but may have developed through the accumulation of simpler building blocks across species, eventually giving rise to a layered structure, characteristic of human language (Levinson and Holler 2014).

In the larger scope of gesture-research in primates, our understanding stems mostly from ape gestures. We now know, as a result of research for over three decades that ape gestures are intentional, dyadic, imperative in nature, developing through ritualization of actions into communicative signals, during an individual’s lifetime (Tomasello and Call 2018). However, phylogenetically conserved signals across ape species also suggest that primate gesture forms may be more regulate through innate mechanisms and characteristics such as flexibility and intentionality must reflect in the way the signals are used (Byrne et al. 2017). In bonnet macaques, we find a repertoire of intentionally produced affiliative, agonistic and play gestures used in dyadic interactions. The gestures are used imperatively, like in the apes, sometimes may be potentially referential (Gupta and Sinha 2016). The lack of individual variation in the gestures used by bonnet macaques leads us to believe that ontogenetic ritualization may not be the mechanism underlying development of macaque gestures. However, we have recently reported a novel manual gesture—hand extension—displayed by some individuals in this macaque population, in the context of requesting food from humans (Deshpande, Gupta, and Sinha 2018). Laidre (2012) reported a similar hand-extension gesture in captive mandrills and hypothesised that it could have developed from the action of reaching through a process of ontogenetic ritualization. There, thus, may be certain gestures in bonnet macaques that are produced in more novel situations which could be learnt socially, while others shape through more innate mechanisms. In summary, our study is a first endeavour to document the gestural repertoire of any wild monkey species, including the exploration of goal-directed intentional use of such signals. Having shown that bonnet macaques use gestures to communicate and having provided the first glimpses of their repertoire and usage, we hope that this study would facilitate a better comprehension of gestures in other non-ape species. More extensive comparative studies of communication across taxa, from the great apes to other primates and beyond, should also enable a more complete tracing of the origin and evolution of language-like markers in the primate lineage.

## Acknowledgements

AS is grateful to the Department of Science and Technology, Government of India, for a research grant under the Cognitive Science Research Initiative (Grant Number SR/CSI/44/2008). SG expresses her deepest gratitude to the National Institute of Advanced Studies for a doctoral fellowship, Kantharaju H C of the Karnataka Forest Department for granting permission for this work, Jagadish M and Sharmi Sen for help during data collection, Kirsty Graham for her comments, suggestions and assurance, Kate Grounds for coding video for inter-observer reliability and Debapriyo Chakraborty for crucial inputs in data analyses and Karthik Davey and Sukanta Das for providing logistical support during fieldwork. Finally, SG and AS are deeply grateful to the reviewers who devoted their time in providing valuable suggestions, which has significantly improved this manuscript in due course of the submission process.

