## Supplementary Material 1 for "Gestural Communication of Wild Bonnet Macaques in the Bandipur National Park, Southern India"

**Ethogram of individual and initiated social behaviours displayed by bonnet macaques in the study troops**

***Individual behaviours***

| **Serial Number** | **Behaviour** | **Codes** | **Notes** |
|---|---|---|---|
| 1 | Alarm Calling | AC | |
| 2 | Cough-Like Calling | IC | Only produced by infants |
| 3 | Movement-Calling | MC | |
| 4 | Lost-Calling | LO | |
| 5 | Feeding on Naturally Available Food | FE | |
| 6 | Feeding on Provisioned Food | FR | |
| 7 | Foraging on Naturally Available Food | FO | |
| 8 | Foraging on Provisioned Food | FP | |
| 9 | Autogrooming | GR | |
| 10 | Bipedal Watching | BW | |
| 11 | Coughing | CI | |
| 12 | Looking | LK | |
| 13 | Lying Down | LD | |
| 14 | Moving | MV | |
| 15 | Playing | PY | |
| 16 | Resting | RS | |
| 17 | Sitting | SN | |
| 18 | Sitting Alertly | SA | |
| 19 | Sneezing | SZ | |
| 20 | Urinating | UR | |
| 21 | Tooth-Picking | TP | |
| 22 | Regurgitating with Subsequent Ingestion | VM | |
| 23 | Watching Out for Other Troops | WT | |
| 24 | Yawning | YW | |
| 25 | Sleeping | ZZ | |

| | | |
|---|---|---|
| 26 | Shaking Branches | SB |
| 27 | Tasting Own Oestrous Material | TE |
| 28 | Rubbing Genitals with Hands | RG |
| 29 | Eating Own Ejaculate | EJ |
| 30 | Missing from the Troop | XX |

***Initiated social behaviours***

| **Serial Number** | **Behaviour** | **Codes** | **Notes** |
|---|---|---|---|
| **Affiliative behaviours** | | | |
| 1 | Allogrooming | AG | |
| 2 | Allogrooming with Lip-Smacking | AS | |
| 3 | Biting Gently | BG | |
| 4 | Biting Another Individual's Infant | BI | |
| 5 | Pulling Close to Oneself | CB | |
| 6 | Approaching, Followed by an Affiliative Interaction | CM | |
| 7 | Cheek-Touching | CT | Discarded from gesture analysis as signaller and receiver were not necessarily visually oriented during display of these behaviours |
| 8 | Raising Eyebrows at Another Individual's Infant | EN | |
| 9 | Following | FW | |
| 10 | Fur-Stroking | FT | Discarded from gesture analysis as signaller and receiver were not necessarily visually oriented during display of these behaviours |
| 11 | Greeting-Grunting | GG | |

| | | | |
|---|---|---|---|
| 12 | Affiliative-Grunting | GU | |
| 13 | Allogrooming Another Individual's Infant | GI | |
| 14 | Grappling | GP | |
| 15 | Huddling | HE | |
| 16 | Handling Another Individual's Infant | HI | |
| 17 | Handling Another Individual's Infant with Lip-Smacking | HL | |
| 18 | Holding any Part of the Body Gently | HO | |
| 19 | Hugging with Lip-Smacking | HS | |
| 20 | Hugging without Lip-Smacking | HU | |
| 21 | Hugging Another Individual's Infant | UI | |
| 22 | Jumping on Another Individual's Back | JB | Only produced by infants and juveniles; sometimes used in the context of play |
| 23 | Licking | LI | Discarded from gesture analysis as signaller and receiver were not necessarily visually oriented during display of these behaviours |
| 24 | Lip-Smacking | LS | |
| 25 | Mouth-to-Body Touching | MB | |
| 26 | Mouth-Sniffing | MF | |
| 27 | Mouth-to-Mouth Touching | MT | |
| 28 | Nibbling | NB | |
| 29 | Nuzzling | NZ | |
| 30 | Nuzzling Another Individual's Infant | NI | |
| 31 | Smelling / Sniffing | SO | Discarded from gesture analysis as signaller and receiver were not necessarily visually oriented during display of these behaviours |

| | | | |
|---|---|---|---|
| 32 | Smelling Another Individual's Infant | OI | Discarded from gesture analysis as signaller and receiver were not necessarily visually oriented during display of these behaviours |
| 33 | Patting | PA | |
| 34 | Pulling Another Individual's Infant | PI | |
| 35 | Raising Eyebrows | RB | |
| 36 | Soliciting Allogrooming with Positive Response | SG | |
| 37 | Soliciting Grooming with Negative Response | SR | |
| 38 | Touching | TO | |
| **Agonistic behaviours** | | | |
| 1 | Aggressive Screaming | AM | |
| 2 | Approaching, Followed by an Aggressive Interaction | AP | |
| 3 | Biting Hard | BH | |
| 4 | Displaying a Bared-Teeth Threat | BT | |
| 5 | Chasing | CH | |
| 6 | Eye-Flashing | EF | |
| 7 | Fleeing | FL | |
| 8 | Fear-Screaming | FS | |
| 9 | Fear-Grimacing | GM | Discarded from gesture analysis as signaller and receiver were not necessarily visually oriented during display of these behaviours |
| 10 | Ground-Slapping | GS | |
| 11 | Threat-Growling | GT | |
| 12 | Holding Down Roughly | HD | |
| 13 | Head-Jerking | HJ | |
| 14 | Head-Jerking at Another Individual's | JI | |

| | | | |
|---|---|---|---|
| | Infant | | |
| 15 | Lunging | LU | |
| 16 | Leaping Away from Another Individual | LW | |
| 17 | Displaying an Open-Mouth Threat | OT | |
| 18 | Punishing Own Infant with Head-Jerks and Bite | PH | |
| 19 | Punishing Own Infant with Head-Jerks and Slap | PP | |
| 20 | Pinching | PN | |
| 21 | Pushing Away | PU | |
| 22 | Pushing Away Another Individual's Infant | EI | |
| 23 | Pulling Another Individual's Body or any Part of it Roughly | PT | |
| 24 | Rushing at Another Individual Aggressively | RA | |
| 25 | Screeching | SE | Usually produced by infants |
| 26 | Slapping | SL | |
| 27 | Slapping Another Individual's Infant | SI | |
| 28 | Spot-Jumping | SJ | Also produced in the context of play |
| 29 | Soliciting Agonistic Support with Positive Response | SS | |
| 30 | Soliciting Agonistic Support with Negative Response | SU | |
| 31 | Staring | ST | |
| 32 | Wagging Tail | WT | Discarded from gesture analysis as signaller and receiver were not necessarily visually oriented during display of these behaviours |
| 33 | Showing Aggression to Humans | AH | |
| **Dominance-subordination behaviours** | | | |

| | | | |
|---|---|---|---|
| 1 | Approaching with the Other Retreating | AR | |
| 2 | Genital Fondling | GF | Also produced in the context of affiliation |
| 3 | Mounting | MO | Discarded from gesture analysis as signaller and receiver were not necessarily visually oriented during display of these behaviours |
| 4 | Mounting with Lip-Smacking | MS | Discarded from gesture analysis as signaller and receiver were not necessarily visually oriented during display of these behaviours |
| 5 | Presenting with Positive Response | PR | |
| 6 | Presenting with Negative Response | PG | |
| 7 | Presenting with Lip-Smacking, with Positive Response | PM | |
| 8 | Presenting with Lip-Smacking, with Negative Response | PX | |
| 9 | Retreating | RE | |
| **Sexual behaviour** | | | |
| 1 | Copulating with Ejaculation | CJ | |
| 2 | Copulating without Ejaculation | CO | |
| 3 | Copulating with Lip-Smacking with Ejaculation | CE | |
| 4 | Copulating with Lip-Smacking without Ejaculation | CK | |
| 5 | Copulatory Lip-Smacking | CS | |
| 6 | Saw-Like Vocalisation During the Last Phase of Copulation or After Copulation | CV | |
| 7 | Gazing | GZ | |
| 8 | Herding | HE | |

| | | | |
|---|---|---|---|
| 9 | Inspecting by Tasting Oestrous Material | IE | Also produced in the context of dominance-subordination.<br>Discarded from gesture analysis as signaller and receiver were not necessarily visually oriented during display of these behaviours. |
| 10 | Inspecting by Smelling | IO | Also produced in the context of dominance-subordination.<br>Discarded from gesture analysis as signaller and receiver were not necessarily visually oriented during display of these behaviours |
| 11 | Inspecting Visually | IS | Also produced in the context of dominance-subordination.<br>Discarded from gesture analysis as signaller and receiver were not necessarily visually oriented during display of these behaviours |
| 12 | Inspecting by Touching | IT | Also produced in the context of dominance-subordination.<br>Discarded from gesture analysis as signaller and receiver were not necessarily visually oriented during display of these behaviours |

| | | | |
|---|---|---|---|
| 13 | Copulatory-Bobbing Laterally | LB | |
| 14 | Copulatory-Bobbing Vertically | UB | |
| 15 | Consorting | SX | |
| **Play behaviours** | | | |
| 1 | Biting in Play | PB | |
| 2 | Chasing in Play | PC | |
| 3 | Retreating in Play | PE | |
| 4 | Lunging in Play | PL | |
| 5 | Displaying an Open-Mouth Threat in Play | PO | |
| 6 | Slapping in Play | PS | |
| 7 | Wrestling in Play | PW | |
| 8 | Rushing at Another Individual in Play | RP | |
| 9 | Leaping Away from Another Individual in Play | LY | |
| **Neutral behaviours** | | | |
| 1 | Avoiding | AV | |
| 2 | Moving Away | MA | |
| 3 | Ignoring | IG | |
| 4 | Coo-Calling | CC | |
| 5 | Lifting Arms to Free Nipples | LF | Only infants produced this behaviour towards their respective mothers |
| 6 | Nipple-Fondling | NF | Only infants produced this behaviour towards their respective mothers |
| 7 | Suckling | SK | Only infants produced this behaviour towards their respective mothers |
| 8 | Observing Intently | OB | |
| 9 | Sleeping Together | SP | |
| 10 | Sitting in Contact or within Half- | SW | |

| | | |
|---|---|---|
| | Metre | |
| 11 | Retreating from Another Individual's Infant | RI |
| **Total number of behaviours = 115** | | |
